# Vitexin alters *Staphylococcus aureus* surface hydrophobicity to interfere with biofilm formation

**DOI:** 10.1101/301473

**Authors:** Manash C. Das, Antu Das, Sourabh Samaddar, Akshay Vishnu Daware, Chinmoy Ghosh, Shukdeb Acharjee, Padmani Sandhu, Junaid Jibran Jawed, Utpal C. De, Subrata Majumdar, Sujoy K. Das Gupta, Yusuf Akhter, Surajit Bhattacharjee

**Affiliations:** Department of Molecular Biology & Bioinformatics, Tripura University, Suryamaninagar, Tripura, 799022, India.; Department of Microbiology, Centenary Campus, Bose Institute, CIT Road, Kolkata 700054, India.; Molecular Stress and Stem Cell Biology Group, School of Biotechnology, KIIT University, Bhubaneswar, Odissa 751024, India.; Centre for Computational Biology and Bioinformatics, School of Life Sciences, Central University of Himachal Pradesh, Shahpur, District-Kangra, Himachal Pradesh 176206, India.; Department of Molecular Medicine, Centenary Campus, Bose Institute, CIT Road, Kolkata 700054, India.; Department of Chemistry, Tripura University, Suryamaninagar, Tripura, 799022, India.; Department of Biotechnology, Babasaheb Bhimrao Ambedkar University, Vidya Vihar, Raebareli Road, Lucknow, Uttar Pradesh, India.

**Keywords:** Keywords Cell surface hydrophobicity, Intercellular adhesion, EPS, Vitexin, Biofilm, *In vivo* biofilm model

## Abstract

Bacterial surface hydrophobicity is one of the determinant biophysical parameters of bacterial aggregation for being networked to form biofilm. Phytoconstituents like vitexin have long been in use for their antibacterial effect. The present work is aimed to characterise the effect of vitexin on *S. aureus* surface hydrophobicity and corresponding aggregation to form biofilm. We have found that vitexin shows minimum inhibitory concentration at 252 μg/ml against *S. aureus.* Vitexin reduces cell surface hydrophobicity and membrane permeability at sub-MIC dose of 126 μg/ml. The *in silico* binding analysis showed higher binding affinity of vitexin with surface proteins of *S. aureus.* Down regulation of *dltA*, *ica*AB and reduction in membrane potential under sub-MIC dose of vitexin, explains reduced *S. aureus* surface hydrophobicity. Vitexin has substantially reduced the intracellular adhesion of planktonic cells to form biofilm through interference of EPS formation, motility and subsequent execution of virulence. This was supported by the observation that vitexin down regulates the expression of *ica*AB and *agr*AC genes of *S. aureus.* In addition, vitexin also found to potentiate antibiofilm activity of sub-MIC dose of gentamicin and azithromycin. Furthermore, CFU count, histological examination of mouse tissue and immunomodulatory study justifies the *in vivo* protective effect of vitexin from *S. aureus* biofilm associated infection. Finally it can be inferred that, vitexin has the ability to modulate *S. aureus* cell surface hydrophobicity which can further interfere biofilm formation of the bacteria.

**Importance:** There has been substantial information known about role of bacterial surface hydrophobicity during attachment of single planktonic bacterial cells to any surface and the subsequent development of mature biofilm. This study presents the effect of flavone phytoconstituent vitexin on modulation of cell surface hydrophobicity in reducing formation of biofilm. Our findings also highlight the ability of vitexin in reducing *in vivo S. aureus* biofilm which will eventually outcompete the corresponding *in vitro* antibiofilm effect. Synergistic effect of vitexin on azithromycin and gentamicin point to a regime where development of drug tolerance may be addressed. Our findings explore one probable way of overcoming drug tolerance through application of vitexin in addressing the issue of *S. aureus* biofilm through modulation of cell surface hydrophobicity.

---

Electrical property of bacterial cell surface plays a key role in bacterial resistance to host effectors. Thus charge modification of cell wall and membrane components becomes very significant (1). In *S. aureus* this charge modification occurs by D-alanyl esterification of teichoic acid in the cell wall, which results in an increased positive charge on the cell (2). The addition of D-alanine esters to teichoic acids is typically mediated by the products of the dlt operon, which encodes four proteins: DltA, a D-alanine: D-alanyl carrier protein ligase; DltB, a D-alanyl transfer protein; DltC, the D-alanyl carrier protein; and DltD, a D-alanine esterase. Incorporation of D-alanine also contributes to the virulence of *Staphylococcus aureus.* Increased expression of *dlt* makes cell surfaces more positively charged which incorporate charge bilayer formation between inside and outside of cell surface. As a result cell exerts more hydrophobicity which in turn favours adhesion of cells by polysaccharide intercellular adhesin (PIA) (3). *In vitro* PIA can be synthesized from UDP-*N*-acetylglucosamine as byproducts of the intercellular adhesion (*ica*) locus. In *S. aureus*, *icaABCD* was shown to mediate cell-cell adhesion and PIA production. It was further demonstrated that *icaA* and *icaD* together mediate the synthesis of sugar oligomers *in vitro*, using UDP-N-acetylglucosamine as a substrate (4).

Biofilm comprise of surface-attached microbial communities encased within a self-produced extracellular matrix and are associated with ~80% of bacterial infections in humans (5). Biofilm formation is thought to require two sequential steps: adhesion of cells to a solid substrate followed by cell-cell adhesion, creating multiple layers of cells (1,5). Intercellular adhesion requires the PIA (4). Biofilm formation from planktonic microorganisms often enhances the pathogenic capability of organisms. Bacterial biofilm associated infections are extremely challenging to treat, as biofilm may become refractory to inhibit by the majority of antibacterial drug used in clinical trial. In addition, biofilm also represent a sanctuary site in which bacteria are physically shielded from attack by the host immune system. Bacterial adherence to the surface of animal cells is an important step in the infection process (6) and hydrophobic interactions are thought to be involved in the adherence of bacteria to host tissues (7,8 and 9). Adherence of bacteria to other bacterial surfaces is affected by the change in interfacial free energy which corresponds to the process of attachment. Therefore, targeting the adherence property of planktonic form of bacteria may develop an effective strategy to prevent the formation of community structure i.e. biofilm. We have previously shown that vitexin, a polyphenolic flavone compound, possess significant antibacterial and antibiofilm property (9) against *Pseudomonas aeruginosa* biofilm. In the present study, we have assayed the involvement of bacterial cell surface hydrophobicity in biofilm formation by *S. aureus.* In that direction, we have explored the probable role of natural flavone like vitexin in reducing *S. aureus* surface hydrophobicity in order to reduce formation of biofilm.

## Results

### Antimicrobial activity of vitexin, azithromycin and gentamicin

The antimicrobial activity of vitexin, azithromycin and gentamicin were studied against *S. aureus.* We have observed that vitexin exhibited highest antimicrobial activity at MIC of 252 μg/ml concentration against the *S. aureus.* The MIC of azithromycin and gentamicin was found to be 110 μg/ml and 5 μg/ml respectively. From the data so obtained, we have selected 126 μg/ml, 106 μg/ml, 86 μg/ml, 66 μg/ml, 46 μg/ml, 26 μg/ml sub-MIC doses of vitexin, 55 μg/ml sub-MIC dose of azithromycin and 2.5 μg/ml sub-MIC dose of gentamicin for all subsequent antibiofilm studies.

Most of the antimicrobials are specific for their mode of action and in this context we tested the killing potential of vitexin alone or in combination using a live-dead staining procedure (see Materials and Methods). To ensure that cells do not die due to the treatments we were intuitive about the extent of PI staining profile to compare the treated cells with the untreated ones. For the experiment *S. aureus* cells were grown to stationary phase (O.D~2.5) and subjected to the proposed treatments with vancomycin as the positive control (Fig. 1A) and untreated as negative control (Fig. 1B). MIC (8 μg/ml) dose of vancomycin was used as described previously (10). All samples with proper controls were analyzed through FACS after 48 hrs of treatment. To our expectation, the sub-MIC dose of vitexin (26 μg/ml) (MVTI) alone (3.6%) (Fig. 1C) or in combination with azithromycin (AZM) (5%) (Fig. 1F) or gentamycin (MGT) (2.7%) (Fig 1G) does not increase PI staining significantly as compared to the control (10.4%) (Fig 1A). Thus, staining profile from FACS data analysis shows that the viability of *S. aureus* cells remains unaltered with the proposed treatment profiles. For better comparison, percentages of PI stained dead cell (Q4) in positive control, negative control and treated samples were presented as bar diagram (Fig. 1H).

**Figure 1:**
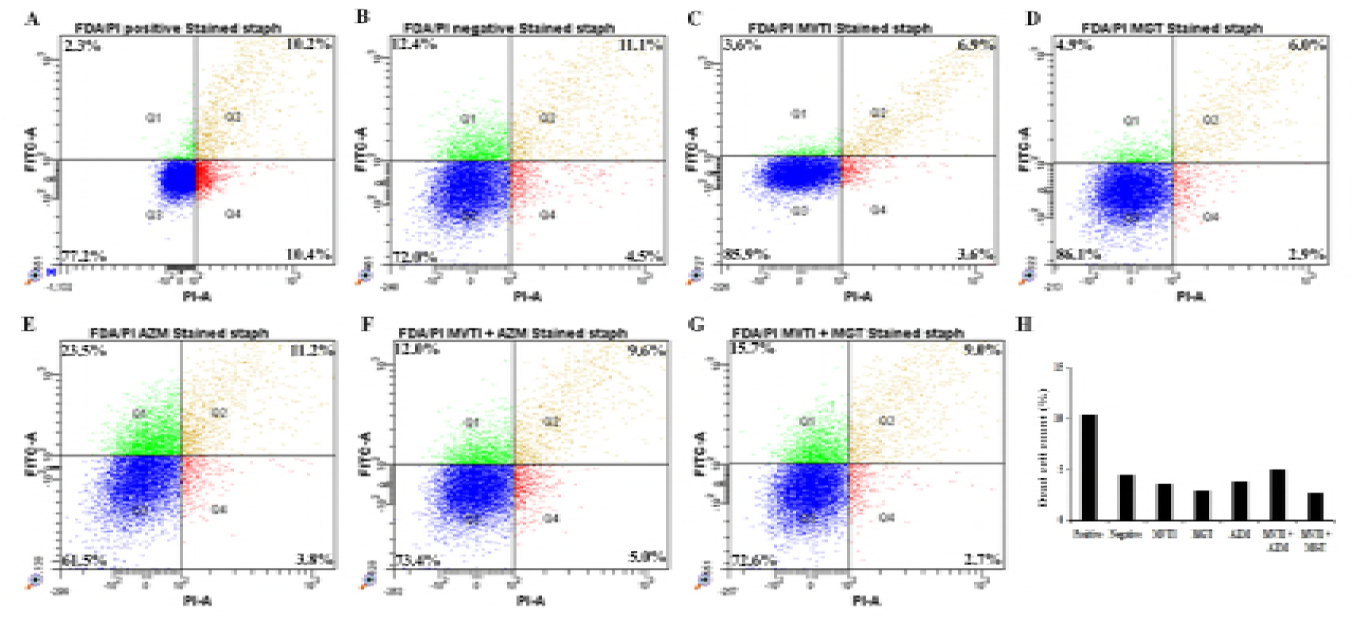
Flow cytometric scatter plot showing live-dead staining profile. The scatter dot-plot is quadrant analyzed using representative colours. The green dots (Q1) represent FDA stained cells, the orange dots (Q2) represent cells stained with FDA and PI, the blue dots (Q3) represent the unstained cells and the red dots represent (Q4) PI stained cells. FDA fluorescence was measured using the FITC-channel and PI stain was measure using PI channel. Vancomycin treated cells were taken as the positive control [A] whereas untreated cells were used as negative control [B]. Vitexin [C], azithromycin [D] and gentamicin [E] treated cells were assayed either alone or in combination as vitexin + azithromycin (MVTI + AZM) [F] or vitexin + gentamicin (MVTI + MGT) [G]. Cell number in the Q3 quadrant indicates the percentage of cells that are PI positive which is a measure of *S. aureus* cell death. Percentage of dead cells from all experimental sets was presented [H]. Box initially at rest on sled sliding across ice.

### Vitexin alters the cell surface hydrophobicity and membrane potential of *S. aureus*

In the present work we have studied the surface hydrophobicity of *S. aureus* after treatment with vitexin alone and in combination. Initially cells were partitioned and cell attachment with acidic solvent and basic solvent was determined. It was found that cell attachments were reduced in basic solvent and increased in acidic solvent. Further, we have observed that hydrophobicity was significantly reduced after administration of vitexin in combination with gentamicin compared to their individual applications [Fig. 2A]. Statistical analysis was performed to determine relation between hydrophobicity (%) and cell attachment (%). The correlation coefficient values for basic solvent was found to be r = 0.9784 and in case of case acidic solvent was r = −0.91496 [Fig. 2A]. We have found that basic solvent slope was negative (−10.284) whereas acidic solvent slope was reverse (8.1297) than that of hydrophobicity (−10.269). This signifies that upon treatment (vitexin alone and in combination) cell surfaces gradually becomes less basic. As a result surface hydrophobicity was reduced which interfere with biofilm formation. All these together signify that cell surface hydrophobicity was the key regulator of cellular adhesion and biofilm formation.

**Figure 2:**
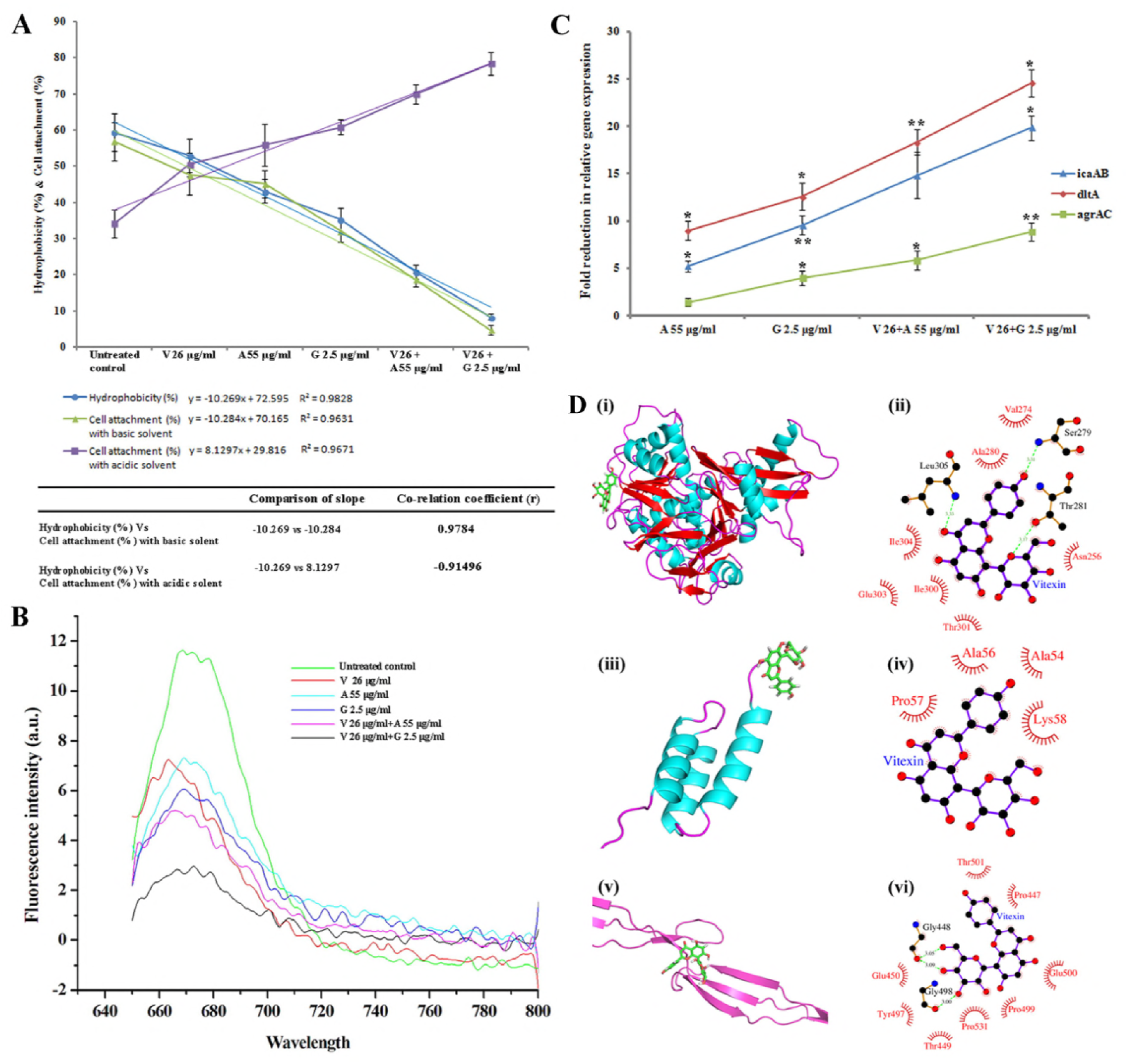
[**A**] Graphical representation and statistical analysis of cell surface hydrophobicity (treated and untreated) in cell attachment (treated and untreated) with acidic and basic solvent. Relationship between these parameters were also analysed through slope of the curve and comparison of correlation coefficient. [**B**] Extent of *S. aureus* membrane depolarisation (treated and untreated) determined through fluorescence intensity of polarisation sensitive dye DiSC_3_. [**C**] Gene expression study of *ica*AB, *dlt*A and *agr*AC gene of *S. aureus.* Changes in gene expression of all treatments were calculated with respect to untreated control taking 16S rRNA as endogenous control. Relative gene expression of A 55 μg/ml, G 2.5 μg/ml, V 26+A 55 μg/ml and V 26+G 2.5 μg/ml were presented with respect to V 26 μg/ml treatment. [**D**] Cartoon representation of protein-ligand complexes with helices coloured in cyan, beta strand in red and coils in magenta colour. Ligand was coloured green and represented in sticks model. Ligplot of protein-ligand complexes are showing interaction of vitexin with protein residues DltA (i, ii), IcaA (iii, iv) and SasG (v, vi). All data were expressed as mean ± SD (n=4 mice per group). *P<0.01, **P<0.001 and **P<0.001 compared with infected mice and calculated through one way ANOVA. Pearson’s Correlation method was used for determining correlation coefficient.

Further to know the effect of these compounds on bacterial cell membrane, we have studied membrane polarisation of all treated samples with respect to untreated control. We have observed that individual treatment with vitexin (26 μg/ml), gentamicin (2.5 μg/ml) and azithromycin (55 μg/ml) membrane surface potential was declined [Fig. 2B]. The combination treatment of vitexin (26 μg/ml) with gentamicin (2.5 μg/ml) significantly increases *S. aureus* membrane potential as compared with treatment of individual compounds [Fig. 2B]. This signifies that vitexin can significantly increase the membrane surface charge and likewise reduce cell-cell adhesion to develop biofilm. The effect of vitexin was found to be potentiated in combination with gentamicin.

### Effect of vitexin (alone and in combination) on biofilm regulatory proteins of *S. aureus.*

To further understand the effect of compounds on biofilm regulatory genes, we have determined the relative mRNA expression of *dltA*, *ica*AB and *agr*AC genes using respective primers [**Table S1**]. Results showed that relative gene expressions of these proteins were significantly reduced where highest fold change was observed in case of vitexin-gentamicin combination treatment [Fig. 2C]. Fold changes of gene expressions with respect to vitexin treatment were calculated by taking 16S rRNA as an endogenous control. Furthermore *in silico* molecular binding affinity study was performed to analyse the effect of vitexin on biofilm associated protein of *S. aureus.* In that direction, we have carried out molecular docking study and attempted to compute the relative binding affinities of vitexin to these proteins. We have carried out possible docking for the proteins encoded by *Ica*, *Dlt*, *Agr* and *Tar* operon. From the molecular docking data, we have observed that vitexin bind into binding pocket of these proteins. For *ica* operon and *dlt* operon only two proteins i.e. IcaA [Fig. 2D (i, ii)] and DltA [Fig. 2D (iii, iv)] were showing affinity for vitexin, out of which IcaA was accommodating vitexin in its native binding pocket while in the case of DltA vitexin was occupied in a different position than that of its native ligand [Fig. 2D]. In the case of SasG [Fig. 2D (v, vi)] protein the vitexin showed a good affinity and bound to the native ligand binding pocket of these proteins. Further, we have energy minimized all vitexin-protein complexes to confirm the stability of complexes and it was also observed that all complexes were having a significantly lower potential energies [Table 1].

**Table 1:** Binding affinity and potential energy values after energy minimization of quorum sensing regulatory and biofilm formation associated proteins of *S. aureus* with a probable inhibitor molecule vitex.

| S.No. | Receptor protein | PDB ID | Native ligand | Potential energy after energy minimization | Autodock binding score* |
| --- | --- | --- | --- | --- | --- |
| 1. | AgrC | 4BXI | Acetate ion | -5.9808762e+05 | -3.9 |
| 2. | AgrA | 4G4K | Glycerol | -5.9459862e+05 | +10.3 |
| 3. | DltA | - | AMP | -1.2131328e+06 | -6.2 |
| 4. | IcaA | - | Beta-D-Glucose | -2.1266348e+06 | -3.7 |
| 5. | SpA | 4NPE | Thiocyanate | -5.9989606e+06 | -3.0 |
| 6. | SasG | 3TIQ | 2-amino-hydroxymethyl-propane-1,3-diol | -1.4015485e+05 | -3.8 |
| 7. | TarL | - | EDT | -1.1858954e+06 | 53.7 |
| 8. | TarK | - | EDT | -1.1773525e+06 | -3.3 |
| 9. | TarO | - | Magnesium ion | -1.1811139e+06 | -4.8 |

\*Autodock gives a binding score indicating the binding affinity measured in kcal/mol. Negative scores indicate high binding affinity whereas positive scores indicate weak binding.

#### Antibiofilm effect of vitexin on *S. aureus.*

Earlier results explain that vitexin reduces *S. aureus* surface hydrophobicity which have significant role on bacterial cell attachment. In that direction, we have tested effect of vitexin (alone and in combination) on biofilm formation by *S. aureus.* Results of crystal violet staining showed that vitexin at sub-MIC doses exerted significant biofilm attenuation against the microorganism and maximum activity was measured at 126 μg/ml dose [Fig. 3A (i)]. It was also observed that antibiofilm activity of vitexin (26 μg/ml) was synergistically potentiated in combination with azithromycin (55 μg/ml) and gentamicin (2.5 μg/ml). The most potent antibiofilm efficacy was observed in combination with gentamicin. Size of bacterial population in biofilm was determined through total extractable protein from the adhered microbial population. Data showed that vitexin treated samples have less extractable protein contents in comparison to the untreated control [Fig. 3A (ii)]. Maximum (69.43%) biofilm attenuation was observed at 126 μg/ml dose of vitexin whereas, azithromycin (55 μg/ml) and gentamicin (2.5 μg/ml) have executed synergistic effect on 26 μg/ml of vitexin in attenuating biofilm total. Consistent with the CV staining assay, protein extraction assay also showed a similar pattern of biofilm attenuation [Fig. 3A (i) and 3A (ii)].

**FIG 3:**
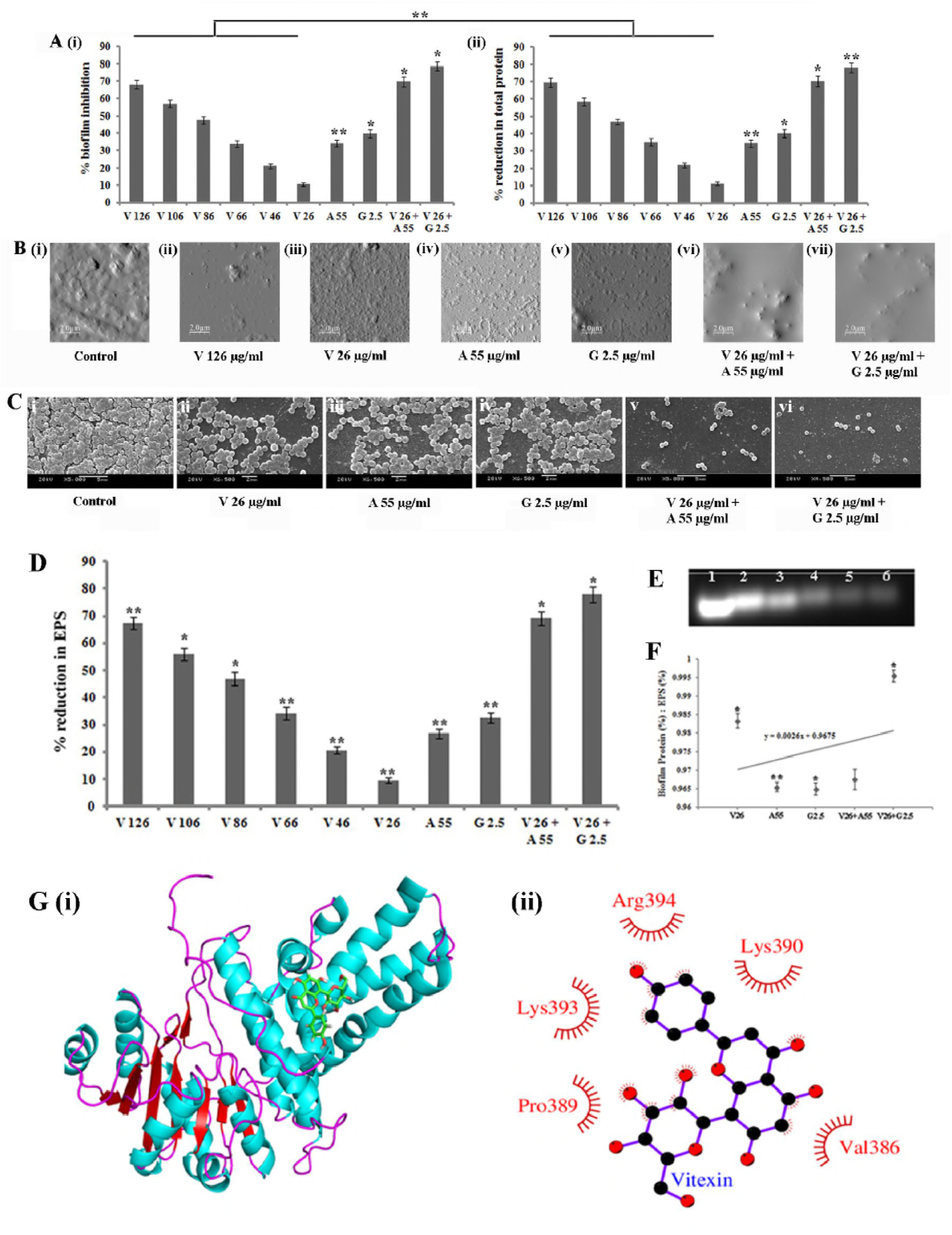
Effect of sub-MIC doses of vitexin and in combination with azithromycin and gentamicin against *S. aureus* on biofilm inhibition [**A (i)**] and inhibition in biofilm total protein [**A (ii)**]. [**B**] Observation of vitexin (alone and in combination) treated and untreated biofilm under atomic force microscope at 2 μm scale. [**C**] Observation of treated *S. aureus* biofilm and bacterial attachment with respect to untreated control through Scanning Electron Microscope. [**D**] Inhibition (percentage) in *S. aureus* EPS formation after treatment with vitexin and in combination with azithromycin and gentamicin with respect to untreated control. [**E**] Agarose gel electrophoresis of eDNA extracted from untreated and treated *S. aureus.* Band intensity of vitexin (2), azithromycin (3), gentamicin (4), vitexin-azithromycin (5) and vitexin-gentamicin (6) were compared with respect to untreated control (1). [**F**] Comparative analysis of modulation of EPS and biofilm protein as an indicator of biofilm inhibition through determination of ratio of biofilm total protein (%) and EPS (%). [**G**] Cartoon representation of protein-ligand complexes with helices coloured in cyan, beta strand in red and coils in magenta colour. Ligand was coloured green and represented in sticks model. Ligplot of protein-ligand complexes are showing interaction of vitexin with protein residues SpA (I, ii). All data were expressed as mean ± SD (n=4 mice per group). *P<0.01, **P<0.001 and **P<0.001 compared with infected mice and calculated through one way ANOVA.

Furthermore, attenuation in biofilm formation by sub-MIC doses of vitexin alone and in combination were further validated by observation under AFM and SEM. AFM snapshots usually showed only surface topology of the cover slip surface from which cell-cell adhesion may not be explained properly. Thus to understand cellular adhesion under high magnification, cover slips from experimental sets were observed under SEM. AFM [Fig. 3B] and SEM [Fig. 3C] results shows that vitexin at sub-MIC doses significantly reduces cellular attachment leading to formation of biofilm. In the control set, untreated cells exert very prominent biofilm over the glass surface. We have also observed that vitexin (26 μg/ml) significantly potentiates the attenuation of cellular adhesion by gentamicin (2.5 μg/ml). Azithromycin (55 μg/ml) also shows moderate biofilm attenuation and cellular adhesion. Taken together, all these results indicate that vitexin at its sub-MIC (126 μg/ml) exhibited moderate biofilm attenuation activity whereas 26 μg/ml concentration of vitexin significantly potentiate the activity of gentamicin to inhibit biofilm formation.

#### Vitexin significantly reduces the *S. aureus* EPS

Biofilm is the cluster of planktonic cells attached by EPS. In the present work, we have quantified the amount of EPS with or without vitexin treatment. We have observed a significant correlation (r = 0.951) between the measured quantity of EPS with antibiofilm activity of vitexin. Vitexin (126 μg/ml) treated samples showed significantly less quantifiable EPS whereas maximum EPS quantities were recorded (9.8% inhibition) in 26 μg/ml dose of vitexin [Fig. 3D]. But after combining sub-MIC doses of azithromycin and gentamicin separately with 26 μg/ml dose of vitexin, EPS quantities were reduced significantly (69.9% and 78.2% inhibition respectively) [Fig. 3D]. In addition to that, EPS DNA (eDNA) were also extracted from all samples (treated and untreated control). Agarose gel electrophoresis showed that eDNA quantity was significantly reduced after treatment with vitexin at 126 μg/ml dose. eDNA quantity was also significantly reduced after vitexin (26 μg/ml)-gentamicin (2.5 μg/ml) combination treatment than their respective individual application [Fig. 3E]. Ratio between biofilm total protein and EPS reveals that with treatment reduction rate of protein was higher than that of EPS but maintains a steady state (slope = 0.0026). But suddenly in vitexin-gentamicin combined treatment, biofilm total protein and EPS ratio becomes significantly high [Fig. 3F]. This signifies that vitexin (26 μg/ml)-gentamicin (2.5 μg/ml) combination treatment have significantly reduced quantity of EPS. All these results validated that vitexin has potent antibiofilm activity at higher sub-MIC doses whereas reduced antibiofilm activity of gentamicin at very low doses was potentiated in combination with sub-MIC dose of vitexin (26 μg/ml). *In silico* binding affinity study of SpA protein with vitexin shows higher binding affinity which validates earlier result of reduction of *S. aureus* EPS after treatment with vitexin [Fig. 3G].

#### Vitexin attenuated *S. aureus* sliding movement and release of bacterial protease

Sliding motility is a kind of bacterial movement which is key regulator of biofilm formation by any bacteria. In the present work, we have observed that vitexin (126 μg/ml) treated cells showed marked reduction in sliding movement compared to the negative control [Fig. 4A (ii)]. It was also observed that sliding motility at vitexin (26 μg/ml)-gentamicin (2.5 μg/ml) treatment [Fig. 4A (vii)] was significantly less than that of their individual treatment. Furthermore, results have also demonstrated that vitexin (126 μg/ml) has executed significant inhibition in protease production by *S. aureus* [Fig. 4B]. It was also observed that lowest dose of vitexin (26 μg/ml) significantly increase the extent of attenuation of proteases by azithromycin and gentamicin. Among these, combination with gentamicin executed higher attenuation than combination with azithromycin.

**Figure 4:**
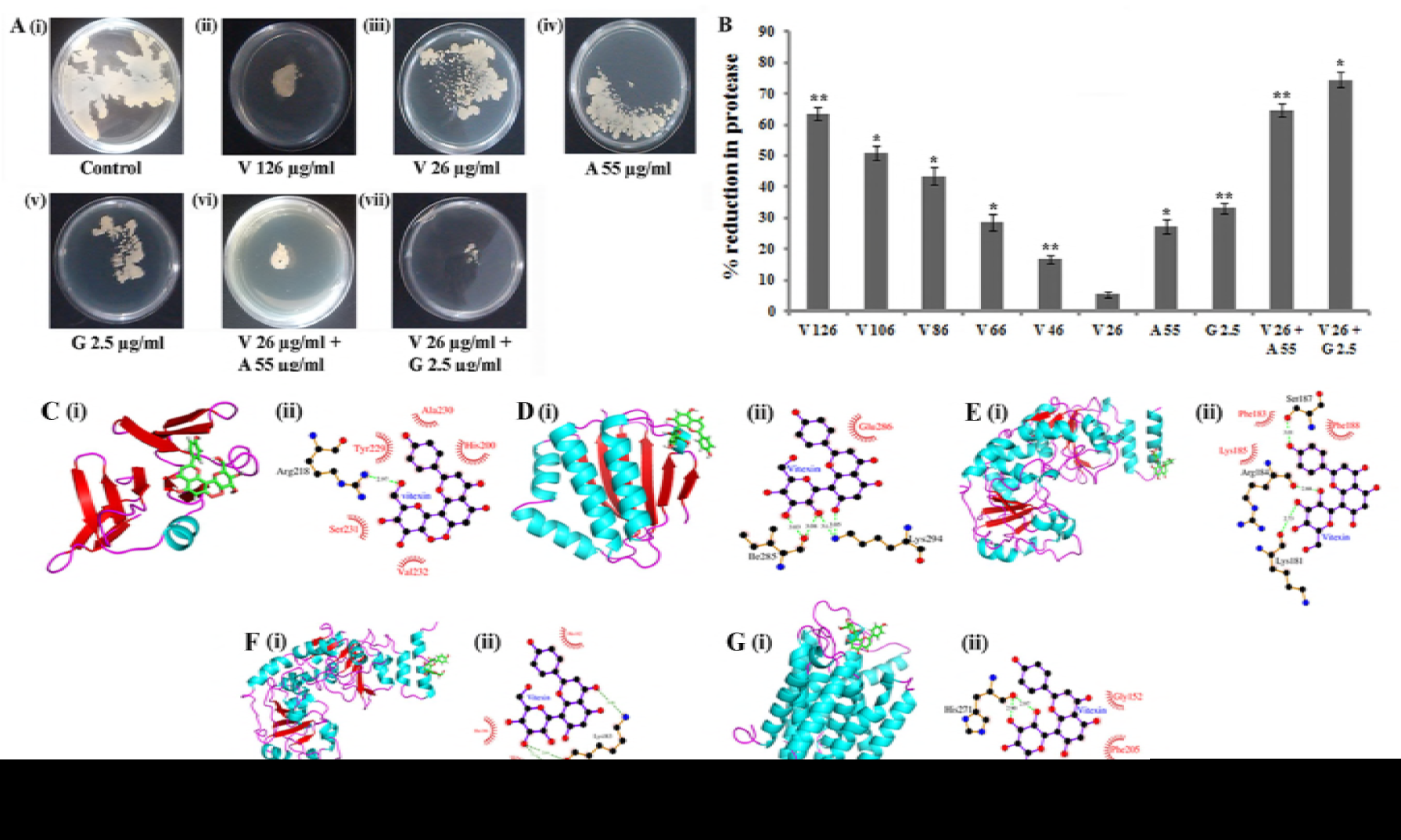
Attenuation of sliding movement [**A**] and protease secretion [**B**] by *S. aureus* after treatment with vitexin (alone, in combination with azithromycin and gentamicin). Cartoon representation of protein-ligand complexes with helices coloured in cyan, beta strand in red and coils in magenta colour. Ligand was coloured green and represented in sticks model. Ligplot of protein-ligand complexes are showing interaction of vitexin with protein residues AgrC [**C**(i)(ii)], AgrA [**D**(i)(ii)], TarL [**E**(i)(ii)], TarK [**F**(i)(ii)] and TarO [**G**(i)(ii)]. All data were expressed as mean ± SD (n=4 mice per group). *P<0.01, **P<0.001 and **P<0.001 compared with infected mice and calculated through one way ANOVA.

To validate results of bacterial motility and virulence, *in silico* molecular docking were performed. We have observed that in *agr* operon AgrC (PDB ID: 4BXI) [Fig. 4C] and AgrA (PDB ID: 4G4K) [Fig. 4D] both were having high binding affinity for vitexin and were occupying the similar binding pocket as that of their native ligand. For biofilm formation associated proteins, TarF and TarO [Fig. 4G] the vitexin binds into the native ligand binding pocket of these proteins, while for TarL [Fig. 4E] and TarK [Fig. 4F] it binds to a different position than the original binding position of the ligand.

#### *In vivo* efficacy of vitexin against catheter-associated infection in a murine model

Furthermore, *in vivo* effects of vitexin (alone and in combination) on attenuation of biofilm was confirmed in mouse model. For that purpose catheter associated biofilm model was developed. At first, effect of these doses on mouse liver and spleen was analysed through histology and subsequently dispersion of catheter associated biofilm was also evaluated through CFU count. The paraffin embedded section of mouse liver and spleen from biofilm infection control groups were compared with treated mouse liver and spleen. It was observed that portal vein and hepatic artery were very much dilated, distribution of hepatocytes were not uniform in liver of infection control mouse. The morphology of the hepatocytes, central vein and hepatic triad varies distinctly among different treatment groups. In mouse treated with vitexin 126 μg/ml, central vein was found regular and the shape of the hepatic lobules was also found to be restored [Fig. 5 (1–6)]. An additive effect of vitexin (26 μg/ml) and gentamycin (2.5 μg/ml) were also observed and it showed highest healing activity where the central vein, hepatic lobule, liver sinusoid, portal triads are found intact and healthy. The regions of splenic nodules, central artery, trabecular vessels, red pulp, and white pulp were identified in control and all the treatment groups [Fig. 5 (7–12)]. Results showed severe structural deformity in the tissue architecture in infection control mouse where it was found to be normalising in vitexin (26 μg/ml)-gentamycin (2.5 μg/ml) treated mouse. In addition to above results, we have determined tissue solidity and roundness through analysis of histological images which depicts that solidity was reduced and roundness was increased with treatment in comparison with untreated control [Fig. 6A]. This implies that tissue architecture was gradually restored after treatment with vitexin alone and in combination.

**Figure 5:**
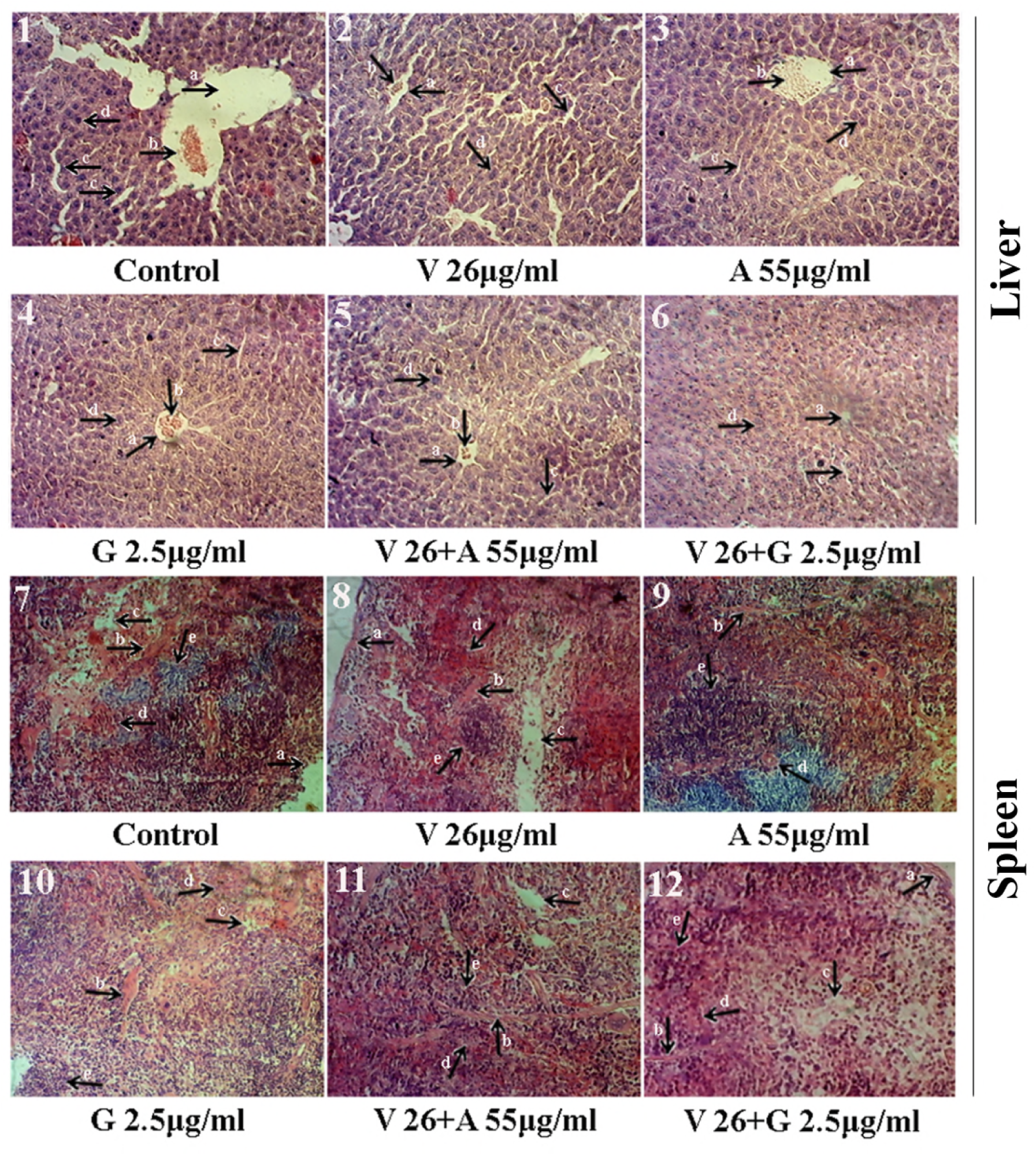
Histopathological examination of mouse (*S. aureus* biofilm model) liver and spleen after all treatments with respect to untreated control [**A**]. In liver a= central vein, b= lymphocytes, c= sinosoids and d= hepatocytes; in spleen a= capsule, b= trabecula, c= central arteriole, d= red pulp and e= white pulp [**A**].

Furthermore, a mouse model of catheter infection was used to evaluate the *in vivo* antibiofilm activity of vitexin alone and in combination with antibiotics. Bacteria were cultivated *in vitro* on implantable catheters and induced to form biofilm in mice. The effects of vitexin (alone and in combination) on catheter associated *in vivo* biofilm are shown in Fig. 6B. Vitexin (1300 μg/Kg-body weight) treatment plate shows 255 cfu/liver in comparison with untreated control where cell count were 436 cfu/liver. This validates the antibiofilm activity of vitexin against catheter associated *in vivo* biofilm form of *S. aureus* infection. This was also observed that combination of vitexin (1300 μg/Kg-body weight) with gentamicin (125 μg/Kg-body weight) treatment shows highest activity (26 cfu/liver) whereas with only gentamicin (125 μg/Kg-body weight) colony count was 205 cfu/liver [Fig. 6B]. Bacterial load after treatment with vitexin (1300 μg/Kg-body weight) and azithromycin (2750 μg/Kg-body weight) treatment was found to be 59 cfu/liver [Fig. 6B]. All these observation firmly validates the *in vivo* antibiofilm activity of vitexin against *S. aureus* biofilm. Results also confirm that sub-MIC dose of vitexin potentiates the activity of sub-MIC dose of gentamicin against *in vivo S. aureus* biofilm.

**Figure 6:**
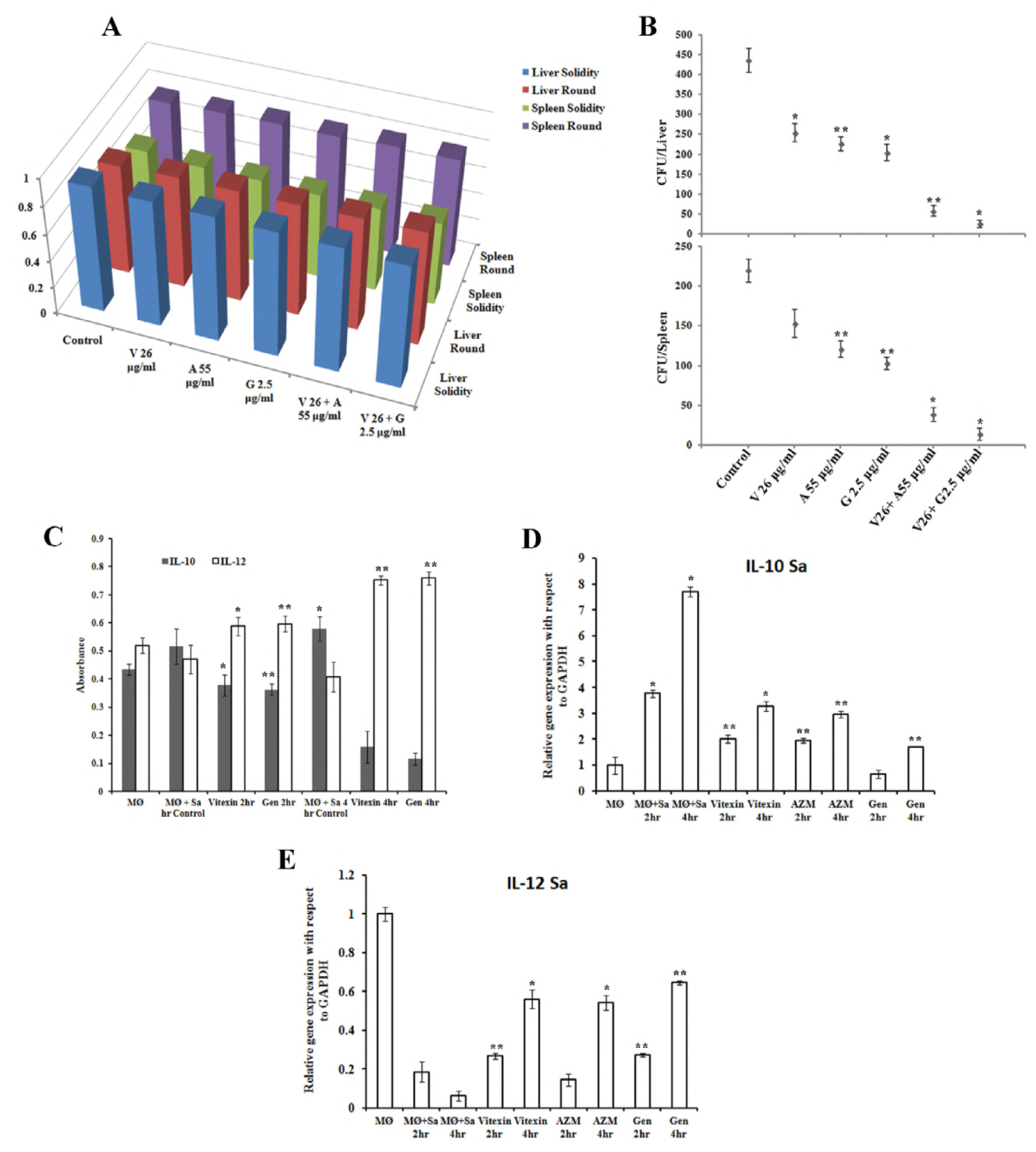
[**A**] *In silico* analysis of tissue (mouse liver and spleen) solidity and roundness of cell through Image-J software. [**B**] Estimation of bacterial load in mouse (biofilm model) liver and spleen determined through CFU count on agar plate. [**C**] Expression of IL-10 and Il-12 cytokines at protein level determined through ELISA. [**D**] IL-10 and [**E**] IL-12 gene expression in RAW macrophages after infection with *S. aureus* biofilm. All data were expressed as mean ± SD (n=4 mice per group). *P<0.01, **P<0.001 and **P<0.001 compared with infected mice and calculated through one way ANOVA.

#### Effect of vitexin on inflammatory response in RAW 264.7 macrophages infected with *S. aureus* biofilm

*In vivo* antibiofilm effect of vitexin was further validated through study of expression profile of inflammatory cytokines. The immune-modulatory effect of vitexin was studied on RAW 264.7 macrophage cells infected with *S. aureus* biofilm. Pro-inflammatory and anti-inflammatory cytokine level were quantified from the culture supernatant of bacteria-macrophage co-culture. In case of *S. aureus* infection it was observed that at 4 hr of infection, Il-10 level was 1.33 fold increased whereas IL-12 level was 1.27.fold decreased at protein level in untreated macrophages [Fig. 6C]. Vitexin treatment was found to reduce IL-10 production and increases the IL-12 level in infected macrophages both at protein [Fig. 6D] and mRNA level [Fig. 6E]. At 4 hr of treatment, IL-10 levels were 3.25 fold and 2.353 fold reduced at protein and mRNA level respectively with respect to infected macrophages. Treatment with gentamicin and azithromycin has found to reduce the IL-10 mRNA expression by 2.595 fold and 4.498 fold respectively [Fig. 6D]. Whereas at 4 hr of vitexin treatment, IL-12 levels were 1.6 fold and 9.33 fold increased at protein [Fig. 6D] and mRNA level [Fig. 6E] respectively with respect to infected macrophages. In case of azithromycin and gentamicin, IL-12 gene expression profiles were 10.67 fold and 9.03 fold elevated [Fig. 6E]. All mRNA fold changes were calculated with respect to untreated infected macrophages and ΔΔCT values were calculated taking GAPDH as endogenous control. Thus cytokine expression study at protein and mRNA level explores that vitexin effectively participate in the immune-modulation through induction of pro-inflammatory and suppression of anti-inflammatory cytokines in macrophages during infection with *S. aureus* biofilm.

## Discussion

Bacterial surface charge and surface property are key regulators toward formation of biofilm. In this context in the present work we have evaluated *S. aureus* surface hydrophobicity after treatment with vitexin. Further we have evaluated *in vitro* and *in vivo* antibiofilm effect of vitexin on biofilm formation by *S. aureus*. Keeping view of increasing drug tolerance, in the present work we have also determined the effect of sub-MIC dose of vitexin on sub-MIC dose of azithromycin and gentamicin.

Microbial biofilm represents a dense association of microorganisms firmly attached to a substratum which creates difficulty in estimating the total number of bacteria in a given biofilm structure (9,11,12). In bacterial population inter-bacterial communication by Quorum Sensing (QS) produces EPS, which forms a network with all the adjacent bacterial colonies to form biofilm (8,13). Furthermore, cell surface hydrophobicity study and membrane depolarisation study explores that treatment with vitexin-gentamicin significantly reduced the cell surface hydrophobicity. Reduction in cell surface hydrophobicity minimizes surface tension of cell surface. *dlt* operon mediates the addition of D-alanine esters to teichoic acids. *dlt* operon encodes four proteins out of which *dlt*A is a D-alanine : D-alanyl carrier protein ligase which is required for successful addition of D-alanine to the cell wall (14). Incorporation of D-alanine into teichoic acid has been demonstrated to increase membrane free charge and resistance of bacteria to antibacterial as well as contribute to the virulence of pathogens. As a result of down regulation of *dlt*A gene cell surface becomes less charged with reduced surface tension. Subsequently, cells treated with vitexin-gentamicin combination utilizes higher quantity of membrane polarisation dye DiSc3, subsequently release very less quantity and shows significantly less fluorescent intensity. In addition to the reduced cell surface hydrophobicity and surface charges, *ica*AB gene was also down regulated. During biofilm formation the adhesion of bacteria to a substrate surface by cell-cell adhesion forms multiple layers of the biofilm (15). This process is associated with the polysaccharide intercellular adhesin (PIA) as a function of *ica* locus (15). It was further demonstrated that *ica*A and *ica*D together mediate the synthesis of sugar oligomers *in vitro*, using UDP-*N*-acetylglucosamine as a substrate for EPS production. Down regulation of *ica*AB leads to reduced intercellular adhesion between cells for colonisation and biofilm formation. In support to this we have observed significantly reduced EPS and eDNA quantity which can reduce bacterial quorum sensing.

The Quorum Sensing (QS) phenomena is activated when bacterial aggregates reach to a threshold of certain population density and is reported to be extensively associated with biofilm formation (16). In order to understand the effect of vitexin and in combination, we have examined the effect of vitexin on QS mediated sliding movement and secretion of proteases by *S. aureus.* In this relation we have observed down regulation of *agr*AC which significantly reduce sliding movement and protease secretion. The accessory gene regulator (*agr*) locus of *Staphylococcus aureus* encodes a two-component signal transduction system that leads to the down-regulation of surface proteins and up-regulation of secreted proteins during *in vitro* growth (16). In essence, *agr*B activity leads to the secretion of the auto inducing pheromone, *agr*D, which binds to and activates the histidine kinase receptor, *agr*C, which subsequently activates the response regulator, *agr*A. The inhibitory activity of these *agr* groups represents a form of bacterial interference that affects virulence gene expression (17). Sliding leads to rapid bacterial translocation and adherence to the surface that promotes efficient colonization of bacterial cells. It was also reported in literature that sliding movement is initiated and functionalises through QS which facilitate bacterial movement from one place to the other. This in turn stimulates bacteria to form biofilm network over the surface. Proteases are products of bacterial metabolism which are hydrolytic in nature that affect the proteins of the host cells (infected tissue), thereby facilitating bacterial invasion and growth (18).

Furthermore, *in vivo* effect of vitexin was also evaluated. Mouse liver and spleen is secondary lymphoid organ which help in metabolizing pathogens and food materials. After insertion of biofilm layered catheter, bacteria will migrate in different organ of the body through blood stream (9,19). As a result bacterial load will rise in liver as well as in spleen. Furthermore, co-culture of bacteria with macrophage also can be used as an efficient tool to study antimicrobial and immunomodulatory effect of any compounds. During infection, host defence counteract the inflammatory response through modulation of expression of pro and anti-inflammatory cytokines. In the present study, we have observed that vitexin provides protection to murine macrophage cell line from *S. aureus* biofilm infection through induction of pro-inflammatory cytokines (20,21).

## Methods

### Flow cytometry and live/dead staining

1 ml of cells (10^6^ cells/ml) (untreated or treated) were stained with the FDA/PI (fluorescein diacetate/propidium iodide) combination stains and kept at 37°C for 2 hr and then washed with PBS and resuspended in the same buffer (22). Cells were analyzed using FACS Aria system (Becton Dickinson, NJ) and data acquisition was done using FACS Diva software.

### Bacterial cell surface hydrophobicity

An aliquot of *S. aureus* culture was inoculated into basal media supplemented with glucose and ammonium sulphate and incubated at 37°C for 2 days. Thereafter, cells were harvested from each experimental set and cell surface hydrophobicity was examined by bacterial adhesion to hydrocarbon (BATH) assay as described previously (23). The formula for measuring cell surface hydrophobicity is as follows:

Cell surface hydrophobicity (%) = 100X{(initial OD-final OD)/initial OD}

### Physicochemical characterization of the cell surface

Hydrophobicity and the Lewis acid/base character of *S. aureus* populations were investigated according to the microbial adhesion to solvents (MATS) method with minor modifications (24).

The percentage of cells associated with each solvent was determined as follows:

Cell attachment affinity (%) = (1-A/A_0_) x 100.

### Membrane depolarization study

The ability of vitexin to depolarize the transmembrane potential of target bacteria was tested by DiSC_3_5-based membrane depolarization assay (25). Cells treated with azithromycin (55 μg/ml) and gentamicin (2.5 μg/ml) was used as positive control samples.

### Gene expression study by real time PCR

RNA from vitexin treated and untreated *S. aureus* was isolated by Trizol. RNA was reverse transcribed to cDNA, gene of interests were amplified using respective set of primers and relative gene expression were quantified by real time PCR using Real-Time PCR Detection System (StepOnePlus, Applied Biosystems) by 2(2DDCt) method. The expression levels of all selected genes were normalized using 16S rRNA as an internal standard (26).

### Antibiofilm activity of vitexin

Interference of biofilm formation upon treatment were performed as the method described in supplementary material for biofilm forming ability of the bacteria. Percentage of biofilm inhibition in all treated wells with respect to untreated controls was determined using the following formula:

Biofilm Inhibition (%) = {(OD of untreated control) − (OD of treated sample) / (OD of untreated control)} X 100.

### Observation of biofilm by atomic force microscopy (AFM) and Scanning Electron Microscopy (SEM)

*S. aureus* was cultured in 35 X 10 mm petridish on the surface of cover slips. After the incubation, cover slips were collected, washed gently with sterile PBS and observed under microscope. In case of atomic force microscopy (Bruker-Innova) films were analysed first at 10 μm scale and gradually up to 2 μm scale at a scanning speed of 1 Hz (27). All images were obtained with a resolution of 512 X 512 pixels.

In case of SEM, cover slips with biofilm were fixed with 2.5% (v/v) glutaraldehyde and the samples were dehydrated with increasing concentrations of tetramethylsilane (TMS) for 2 min each. The samples were stored in vacuum until use. Prior to analysis by SEM [JSM-6360 (JEOL)] samples were subjected to gold sputtering (JEOL JFC 1100E Ion sputtering device). Images were captured from 20 different fields from a single cover slip (28).

### Histopathology of mouse liver and spleen

Livers from mice were fixed in 10% neutral buffered formalin solution, dehydrated in graded alcohol and embedded in paraffin. Paraffin sections of 3-4 micron thickness were obtained, mounted on glass slides and counterstained with hematoxylin and eosin for light microscopic analyses (29). Histochemical analysis of tissues of treated mice with respect to untreated control was done using Image J software.

### Immunomodulatory study of biofilm macrophage co-culture

The bactericidal effect of vitexin on macrophages after infection with *S. aureus* biofilm was performed as described previously by Auriche et al. 2010 with minor modifications. Following that cytokine protein expression were quantitated through ELISA and gene expressions were through qPCR (30).

### Statistical Analysis

All experiments were performed in triplicate. Data were presented as mean ± standard error. All analysis was performed with Graph Pad Prism (version 6.0) software. Significance level was determined by using One way ANOVA and mentioned as *P* value < 0.01 (*), *P* value <0.001 (**) and *P* value <0.0001 (***).

## Acknowledgements

Authors are thankful to Department of Chemistry, Tripura University and Bose Institute for extending their instrumental facility. Authors would also like to thank Dr. Syed Arshad Hussain, Dept. of Physics, Tripura University for instrumental support. Authors also like to acknowledge Dr. Avik Sarkar, Dept. of Molecular Biology & Bioinformatics, Tripura University, India for their effort in finalising the manuscript. Research in Dr. S. Bhattacharjee’s lab is supported by Govt. of India extramural research funds from ICMR and DBT. Research in Dr. Y. Akhter’s lab is supported by Govt. of India extramural research funds from UGC and SERB-DST.

